# PRC2 Restricts Malignant Peripheral Nerve Sheath Tumorigenesis in a Genetically Engineered Mouse Model of MPNST

**DOI:** 10.1101/2025.11.12.687881

**Authors:** Woo Hyun Cho, Amish J. Patel, Makzhuna Khudoynazarova, Sarah Warda, Juan Yan, Dan Li, Dana Schoeps, Eve Fishinevich, Mohini R. Pachai, Cindy J. Lee, Zoe Jacobs, Kae Kristoff, Jessica J. Sher, Cristina R. Antonescu, Yu Chen, Ping Chi

**Author notes:** **Corresponding authors:** Dr. Yu Chen Memorial Sloan Kettering Cancer Center, 1275 York Avenue, New York, NY 10065.; and Dr. Ping Chi, Memorial Sloan Kettering Cancer Center, 1275 York Avenue, New York, NY 10065. These authors contributed equally to this work (Woo Hyun Cho, Amish J. Patel).

## Abstract

Polycomb Repressive Complex 2 (PRC2), which normally regulates transcriptional silencing, chromatin compaction, and stem cell biology, has both oncogenic and tumor suppressor roles in cancer development depending on tumor type. Malignant peripheral nerve sheath tumor (MPNST), characterized by *NF1, CDKN2A* and PRC2 loss, is an aggressive subtype of sarcoma with poor prognosis and no effective therapy. In high-grade human MPNSTs, inactivating mutations in PRC2 core components *SUZ12* or *EED* are prevalent and contributes to oncogenic transformation and progression of MPNST. How PRC2 inactivation contributes to MPNST pathogenesis, however, remains incompletely understood. Here we show that genetic inactivation of *Eed* in addition to *Nf1* and *Cdkn2a* in the Schwann-progenitor lineage leads to widespread tumorigenesis within the sciatic nerve compartment of mice. In contrast, loss of *Nf1* and *Cdkn2a* is insufficient to drive tumorigenesis in the sciatic nerve but leads to MPNST development in other anatomic locations with a longer latency. Single-nucleus multiome sequencing of the sciatic nerves revealed that PRC2-loss reprograms *Nf1*/*Cdkn2a*-deficient Schwann-lineage cells toward a dedifferentiated, neural crest stem cell-like state that resembles the transcriptomic signatures of human PRC2-loss MPNST. Together, these findings suggest a context-dependent tumor suppressive role for PRC2 within the sciatic nerve and establish a novel mouse model that recapitulates human PRC2-loss MPNST.

**SIGNIFICANCE:** We present a novel genetically engineered mouse model that faithfully recapitulates human PRC2-loss MPNST, enabling mechanistic and preclinical studies of malignant transformation in the context of PRC2 loss.

## INTRODUCTION

MPNST is an aggressive soft tissue sarcoma with a dismal prognosis due to challenges in local surgical resection and a lack of effective systemic therapies (1). They clinically manifest in different patient settings: Neurofibromatosis type I (NF1)-associated (45%), sporadic (45%), and radiotherapy (RT)-associated (10%), but share recurrent biallelic inactivating mutations in three tumor suppressor pathways: *NF1, CDKN2A*, and (*SUZ12* or *EED*) (2-4).

NF1 is a tumor predisposition genetic disorder impacting 1 in 3,000 people worldwide due to germline heterozygous loss-of-function mutations in the *NF1* tumor suppressor gene that encodes a negative regulator of RAS (5). About 30-50% of NF1 patients develop plexiform neurofibromas (pNFs), which are benign peripheral nerve sheath tumors (PNSTs) that have *NF1* loss of heterozygosity (*LOH*) (6). Consistent with this, deletion of *Nf1* in the Schwann cell lineage can lead to pNF in mice (7-9). A subset of human pNFs can progress to atypical neurofibroma (also known as atypical neurofibromatous neoplasms of uncertain biologic potential - ANNUBP), which typically have 9p21 genomic deletions centering on the *CDKN2A* tumor suppressor gene (10-12). Atypical neurofibromas (ANFs) have nuclear atypia and increased cellularity without necrosis (11). Both pNFs and ANFs can undergo malignant transformation into MPNST (10-12). Interestingly, loss of both *Nf1* and *Cdkn2a* is associated with progression to ANF in 30-40% of mouse tumors, and ∼40% of these tumors progress to MPNST when transplanted into new mice (13, 14). In humans, up to 70% of NF1-associated MPNSTs, but not pNF nor ANF, have homozygous loss-of-function mutations in either *SUZ12* or *EED*, which are members of the Polycomb Repressive Complex 2 (PRC2) (3). PRC2 facilitates transcriptional silencing and chromatin compaction through H3K27me2/3, and its inactivation leads to global H3K27me2/3 loss; hence, the loss of H3K27me2/3 by immunohistochemistry has been widely adapted as a diagnostic biomarker of PRC2 loss and an indicator of malignant transformation of neurofibroma to MPNST (3, 15, 16). It remains unclear how PRC2 inactivating mutations might contribute to MPNST pathogenesis.

Here, we investigated the impact of conditional deletion of *Nf1, Cdkn2a*, and *Eed* in Schwann cell lineage on peripheral nerve sheath tumor pathogenesis in mice. We found that deletion of *Nf1* and *Cdkn2a* led to increased cellularity in the peripheral nervous system and the development of typically one to two MPNST tumors per mouse in various locations without loss of PRC2 function. In contrast, co-deletion of *Eed* with *Nf1* and *Cdkn2a* consistently led to debilitating hindlimb paralysis associated with gross enlargement of lumbar nerves (from paraspinal to sciatic) that histologically resemble hyper-cellular ANFs and low-grade MPNST. These observations indicate that co-deletion of *Nf1* and *Cdkn2a* is required for the development of murine MPNST, but additional tumor suppressive mechanisms impede the efficiency of MPNST. In this case, PRC2 represents an epigenetic tumor suppressor in which its inactivation via *Eed* deletion accelerates *Nf1*/*Cdkn2a* loss driven nerve sheath tumorigenesis broadly in the peripheral nervous system. To dissect the molecular basis of this transformation, we performed Multiome single nucleus (sn) ATAC-seq and snRNA-seq of age-matched sciatic nerves from mice with different genotypes (wild-type control [WT], *Nf1/Cdkn2a*^*-/*-^ [PNC], and *Nf1/Cdkn2a/Eed*^*-/-*^ [PNCE]). These analyses revealed the emergence of a distinct tumor cluster characterized by loss of Schwann cell identity, activation of developmental and EMT-associated transcriptional programs, and enrichment of PRC2-loss signatures observed in human MPNST. Upon transplantation into immunocompetent mice, PRC2-loss sciatic nerves gave rise to high-grade MPNST, showing that these lesions progress to advanced malignancy with time. This mouse model is one of the first to use a conditional genetically engineered mouse model (GEMM) to investigate the contribution of PRC2-loss, along with *Nf1*/*Cdkn2a* co-deletion, in NF1-associated MPNST development and progression, and provides a valuable model for molecular and therapeutic intervention studies in immunocompetent mice.

## MATERIALS AND METHODS

### Animal studies

Animal experiments were carried out in accordance with protocols approved by the MSKCC Institutional Animal Care and Use Committee (IACUC) and were in compliance with relevant ethical regulations regarding animal research. Plp-CreERT mice were obtained from The Jackson Laboratory (strain# 005975, RRID:IMSR_JAX:005975)(17). *Nf1* flox mice were obtained from The Jackson Laboratory (strain# 017639, RRID:IMSR_JAX:017639) (18). *Cdkn2a* flox mice were obtained from The Jackson Laboratory (Cdkn2a^tm4Rdp^, MGI:2687203) (19). Scott Armstrong provided *Eed* flox mice (20, 21). Rosa26-lox-stop-lox-EYFP mice were obtained from The Jackson Laboratory (strain# 006148) (22). Tamoxifen (Toronto Research Chemicals) was dissolved in corn oil at 20 mg/mL final concentration. CreERT activity was induced in neonatal mice (0 to 3 days old) with a single subcutaneous administration of 0.4 mg tamoxifen.

### Immunohistochemistry

All tissues were fixed overnight in 4% paraformaldehyde (2 hours for nerve tissues) followed by thrice washes with 1x PBS, placed in 70% ethanol and sent to Histoserv, Inc for paraffin embedding. Hematoxylin and eosin (H&E) staining was performed by Histoserv, Inc. Immunohistochemistry of all tissues was carried out on a Ventana BenchMark ULTRA Automated Stainer (RRID:SCR_021254). The following primary antibodies were used in this study: Ki67 (ab16667, Abcam, 1:200), H3K27me3 Lys27 (9733, Cell Signaling Technology, 1:100), S100B (ab52642, Abcam, 1:10,000), ROBO2 (45568, Cell Signaling Technology, 1:100). Slides were scanned into digital files by the MSKCC Molecular Cytology core facility using a Mirax scanner. Images were captured from digital slides via CaseViewer software (3DHISTECH).

### PCR genotyping of mouse strains

DNA from tissues was quickly extracted from tissues by incubation at 98°C for 1 hour in 25 mM NaOH/0.2 mM EDTA followed by the addition of an equal volume of 40 mM Tris-HCl (pH 5.5). PCR was carried out with genotyping primers listed in **Supplementary Table 1**. The resulting PCR products were separated by size in an ethidium bromide-stained agarose gel by electrophoresis. Gels were placed into an Alpha Imager HP gel imaging system (Alpha Innotech) for UV exposure and subsequent imaging.

### Nuclei extraction for single-nucleus 10x Multiome ATAC + Gene Expression assay

Sciatic nerves were collected from 15-week-old WT, PNC, and PNCE mice and snap-frozen in liquid nitrogen for storage. Nuclei were isolated from frozen tissues using the Singulator S100 (S2 Genomics) according to the published protocol (DOI: 10.17504/protocols.io.j8nlkkmq6l5r/v1). Briefly, each frozen sciatic nerve was homogenized and processed for nuclei extraction, washed in sucrose-supplemented buffer, and fluorescence-activated cell sorted (FACS) to enrich for 7-AAD-positive nuclei. Purified nuclei were then used for 10x Genomics Chromium Next GEM Single Cell Multiome ATAC + Gene Expression kit following the manufacturer’s instructions.

### Single-nucleus Multiome data analysis

Reads obtained from the 10x Genomics single-nucleus Multiome platform were mapped to mouse genome (mm10) using the “cellranger-arc” software (10x Genomics). The RNA reads were filtered using scCB2 package and doublets were detected and filtered out using the doublet detection package. The resulting fragment files and count matrices were analyzed using Seurat version 5.2.1 and ArchR 1.0.3, and implemented in R version 4.4.2. Nuclei were retained for downstream analysis if they met the following quality-control thresholds: RNA counts between 1,000 and 20,172; detected genes between 500 and 6,650; mitochondrial RNA content <5% of total transcripts; and transcription start site (TSS) enrichment ≥4. Standard Seurat and ArchR workflows were used for data normalization, dimensionality reduction, and integration of RNA and ATAC modalities. snRNA-seq and snATAC-seq datasets were analyzed independently. For gene expression analysis, 20 principal components were used for Seurat’s FindNeighbors function, and clustering was performed using Leiden algorithm with a resolution of 1 in FindClusters function. Clusters detected exclusively in single biological replicates were excluded, and the remaining cells were re-clustered using the same parameters. To project human single-cell MPNST datasets onto mouse data, human genes were first converted to mouse orthologs, and both datasets were subset to shared genes. Cross-species mapping was then performed using Seurat’s FindTransferAnchors and TransferData functions. For chromatin accessibility analysis, doublets were removed using ArchR’s built-in doublet detection module. Peaks were identified using MACS2 v2.2.7.1 on pseudo-bulk replicates for each cluster, with default parameters.

## RESULTS

### PRC2-loss leads to sciatic nerve enlargement in PNCE mice

To investigate the impact of *Nf1, Cdkn2a*, and PRC2 in nerve sheath tumorigenesis, we first obtained a tamoxifen-inducible Plp-CreERT mouse line previously shown to express Cre in Schwann cells upon tamoxifen administration (8, 17). We then bred Plp-CreERT mice with *Nf1, Cdkn2a, Eed* mice until we obtained three cohorts of genetically engineered mouse models: 1) Plp-CreERT: *Nf1*^fl/fl^ *Cdkn2a*^fl/fl^ (PNC) 2) Plp-CreERT: *Nf1*^fl/fl^ *Cdkn2a*^fl/fl^ *Eed*^fl/fl^ (PNCE) 3) *Nf1*^fl/fl^ *Cdkn2a*^fl/fl^ *Eed*^fl/fl^ (WT). Tamoxifen was administered to neonates from all three genotypes to induce Cre-mediated recombination, and animals were aged to monitor for any phenotype and survival (**Fig. 1A, S1A-C**).

**Figure 1.**
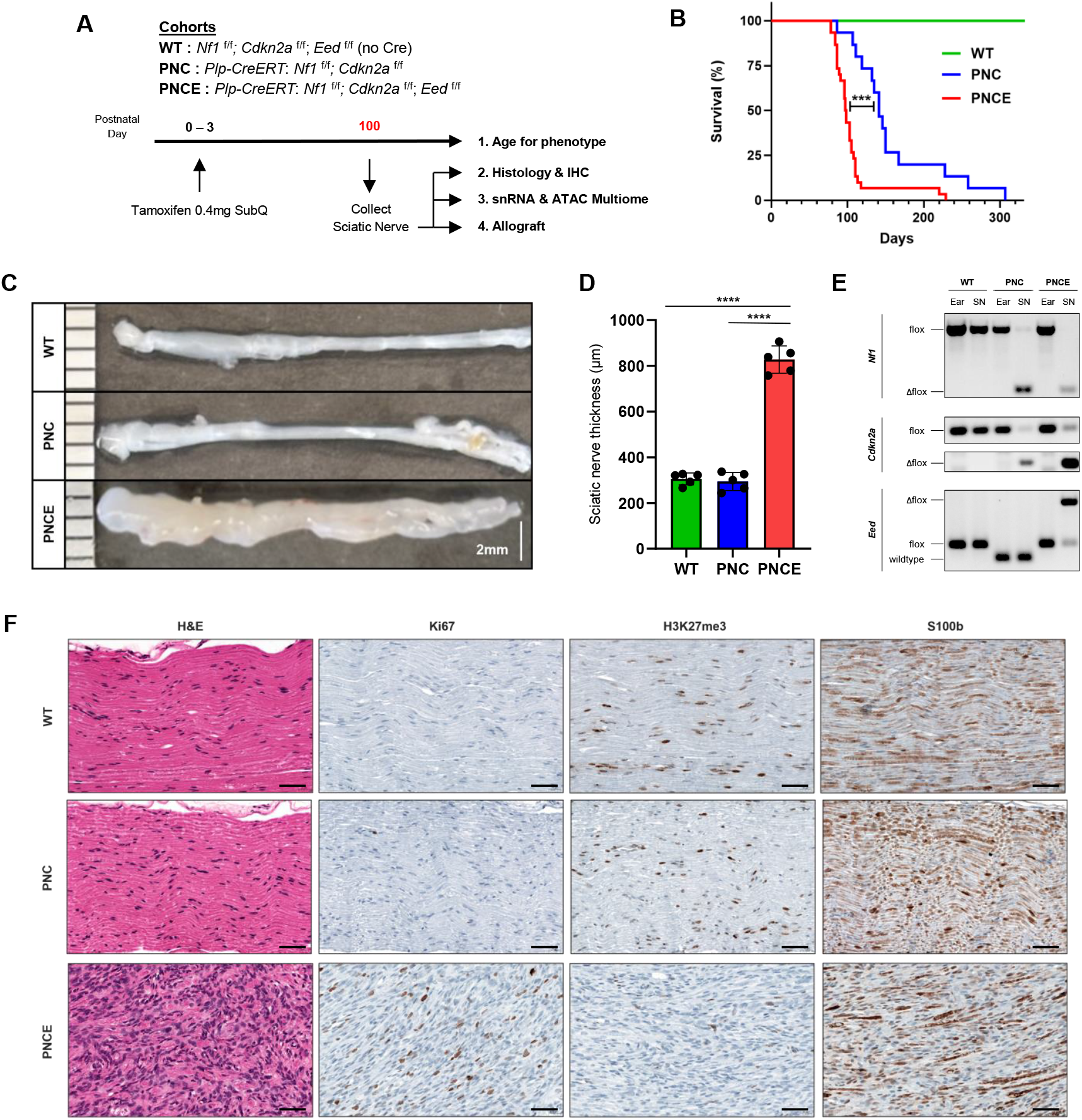
PRC2 loss leads to sciatic nerve enlargement in PNCE mice. **A**. Schematic of genetically engineered mouse models and experiment setup. Neonatal pups (P0-P3) received 0.4 mg tamoxifen to induce Plp-CreERT mediated recombination. Mice were aged to assess phenotype and sciatic nerves were collected at ∼100 days (PNCE) or 15 weeks for histology, single-nucleus multiome, and allograft studies. **B**. Kaplan-Meier survival curves of each mouse cohort (PNC, n=15; PNCE, n=30). Statistical significance was determined by log-rank (Mantel-Cox) test (***, p < 0.001). **C**. Representative gross images of sciatic nerves from each cohort. **D**. Quantification of sciatic nerve thickness measured from hematoxylin and eosin (H&E)-stained sections. Each point represents the mean of three independent measurements per nerve. Data are shown as mean ± SD and analyzed by two-tailed t test (**** p < 0.0001). **E**. PCR genotyping of sciatic nerve samples confirming Cre-mediated deletion of *Nf1, Cdkn2a*, and *Eed*. **F**. Representative H&E and IHC staining of sciatic nerves from 15-week old WT, PNC, and PNCE mice following neonatal tamoxifen administration. Scale bar, 50μm.

At approximately 100 days postnatal, PNCE mice developed severe bilateral hindlimb paralysis and had to be euthanized per IACUC standards. PNC mice remained healthy at this age with no visible symptoms of paralysis (**Fig. 1B**). However, when aged for longer until approximately 145 days, PNC mice had to be euthanized as they developed tumors at various locations (descending from paraspinal to lumbar sciatic region, cervical, and head) (**Fig. S1A**).

Histologically, the PNC tumors were densely packed with spindle-shape cells that have a high nuclear to cytoplasmic ratio, fascicle and storiform structures, with frequent Ki-67-positive cells, and strong H3K27me3 staining, similar to human MPNSTs with intact PRC2 (**Fig. S1D**). This was consistent with previously reported GEMM with *Dhh*-Cre driven deletion of *Nf1* and *Cdkn2a* that develops high grade genetically engineered mouse (GEM)-PNSTs similar to human MPNST by 5 months of age (13).

Contrary to our initial hypothesis, additional deletion of *Eed* in PNCE mice did not accelerate the growth of subcutaneous GEM-PNSTs as seen in PNC mice. No PNCE mice had such discrete lesions at the time of sacrifice. Instead, at necropsy, PNCE mice showed dramatically enlarged sciatic nerves, a feature absent in age-matched WT or PNC mice (**Fig. 1C, 1D**). We confirmed by PCR genotyping that the sciatic nerves from each group had the expected deletions of either *Nf1, Cdkn2a*, and *Eed* (**Fig. 1E**). Histologically, PNCE sciatic nerves at approximately 100 days postnatal were hypercellular with nuclear atypia, Ki67 positivity, and loss of H3K27me3, consistent with low-grade MPNST (**Fig. S2A, Fig. 1F**). In contrast, age-matched PNC sciatic nerves resembled WT sciatic nerves and retained H3K27me3. In addition, PNCE sciatic nerves had multiple areas with loss of S100B immunostainings, indicating loss of Schwann-lineage identity, whereas the WT and PNC sciatic nerves exhibited prevalent and diffuse S100B immunostaining. Because PNCE mice were required to be euthanized due to hindlimb paralysis, these findings likely represent an early stage of malignant transformation driven by PRC2-loss, which will likely progress to high-grade MPNST if time allows. Together, these data show that additional loss of *Eed* in *Nf1*/*Cdkn2a*-deficient Schwann cells is sufficient to initiate low-grade MPNST.

### Single-nucleus RNA and ATAC sequencing identify a tumor cluster in the PNCE sciatic nerve

To better understand the molecular basis of these phenotypic differences, we performed Multiome snATAC-seq and snRNA-seq of the sciatic nerves harvested from WT, PNC, and PNCE mice 15-week post neonatal tamoxifen administration. We pooled sciatic nerves from 2 to 3 mice for each replicate, and after quality control, analyzed a total of 30,884 nuclei from WT (n=2), PNC (n=3), PNCE (n=3) sciatic nerves (**Fig. S3A**). We visualized the transcriptomes and chromatin accessibilities of these nuclei on uniform manifold approximation and projection (UMAP) and used Leiden clustering to identify clusters within the sciatic nerve. Cell types were annotated using previously reported single-cell datasets of mouse sciatic nerve as well as known canonical marker genes (23, 24). This allowed us to identify a novel tumor cluster unique to the PNCE group in both the RNA UMAP and ATAC UMAP (**Fig. 2A, 2B**). Marker gene analysis with a threshold of genes expressed in 70% of cells in the tumor cluster identified *Robo2, Cdh11, Col11a1, Lsamp*, and *Fmo1* as tumor-associated genes (**Fig. 2C, 2F**). Notably, *Robo2, Col11a1*, and *Lsamp* were also among the genes significantly upregulated in PRC2-loss compared with PRC2-wild-type MPNSTs in our previously published human RNA-seq dataset (25).IHC of the sciatic nerves confirmed the increase in ROBO2 staining only in the PNCE sciatic nerve (**Fig. 2G**). When the composition of cell types was compared, non-myelinating Schwann cells (nmSC) were the most expanded cluster in the PNC sciatic nerve compared to the WT sciatic nerve (2.2% in WT and 43.9% in PNC) (**Fig. 2D**). Composition of myelinating Schwann cells (mSC) only showed minor changes (5.5% in WT and 6.9% in PNC), suggesting that Plp-CreERT driven deletion of *Nf1* and *Cdkn2a* preferentially affects nmSCs to proliferate. In the PNCE group, tumor cells were the majority cell type, comprising 61% of all cells in the sciatic nerve. Notably, we identified a unique cluster of cells in the PNCE sciatic nerve that expressed both Schwann cell and Tumor markers, which we termed “SC_Tumor”. SC_Tumor cluster expressed lower levels of Schwann cell marker genes and higher levels of Tumor marker genes than Schwann cell cluster, likely representing cells undergoing lineage transition from Schwann cells to tumor cells (**Fig. 2E**). These findings suggested that the loss of *Nf1* and *Cdkn2a* drives the proliferation of nmSCs while preserving Schwann cell identity, but additional *Eed* loss releases their commitment to Schwann lineage, consistent with PRC2’s role in maintaining differentiation.

**Figure 2.**
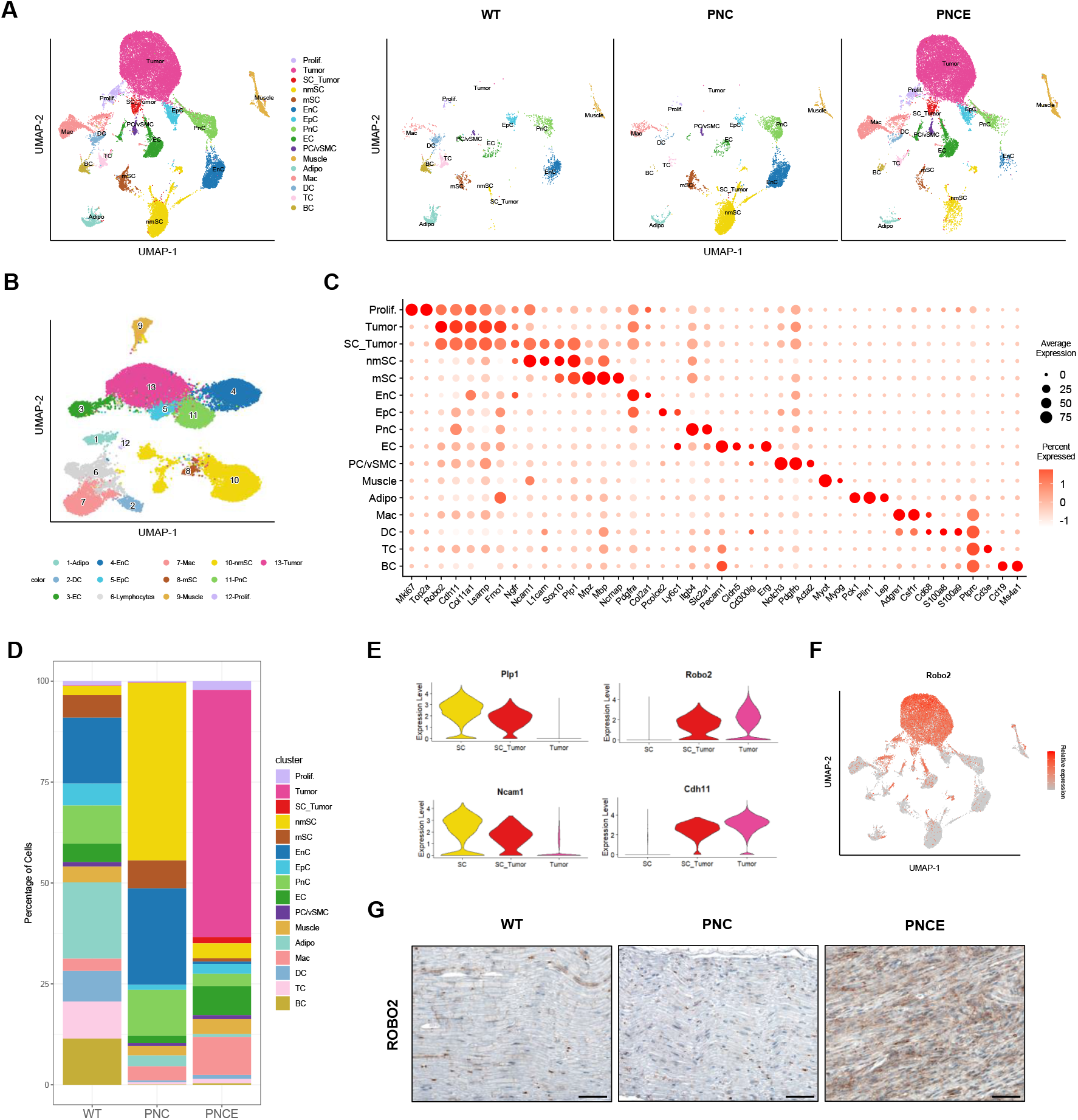
Single-nucleus RNA and ATAC sequencing identify a tumor cluster in the PNCE sciatic nerve. **A**. UMAP plot of snRNA-seq data showing cell clusters in pooled WT, PNC, and PNCE sciatic nerves (left), and separated by genotype (right). **B**. UMAP plot of single-nucleus ATAC-seq (snATAC-seq) data showing corresponding chromatin accessibility-based clusters. **C**. Dot plot of scaled expression of canonical marker genes differentiating major cell types in the mouse sciatic nerve. **D**. Proportion of cell types in each mouse cohort. **E**. Violin plots showing expression of Schwann cell markers (*Plp1, Ncam1*) and tumor cell markers (*Robo2, Cdh11*) across the Schwann cell, SC_Tumor, and Tumor clusters. **F**. Scaled expression of *Robo2* visualized on the snRNA-seq UMAP. **G**. IHC validation of ROBO2 expression in WT, PNC, and PNCE nerves. Scale bar, 50μm.

### PNCE tumor cluster resembles human PRC2-loss MPNST and expresses neural crest stem cell (NCSC)-associated transcription factors

Based on our hypothesis that PNC nmSC cluster represents an earlier stage in tumor progression characterized by *Nf1* and *Cdkn2a* loss alone, we compared PNCE Tumor cluster with PNC nmSC cluster to define the transcriptional programs associated with PRC2 loss. Gene Set Enrichment Analysis (GSEA) revealed strong enrichment of custom gene set comprising top 300 genes upregulated in PRC2-loss versus PRC2-wildtype (WT) human MPNST (**Fig. 3A**) (25). When per-cell module scores were calculated based on the same gene set, we observed that PNCE Tumor cluster had a significantly higher PRC2 loss module score than PNC nmSC cluster (**Fig. 3B**). In contrast, PNC nmSC cluster scored higher when the top 300 genes upregulated in PRC2-WT human MPNST were used (**Fig. 3C**).

**Figure 3.**
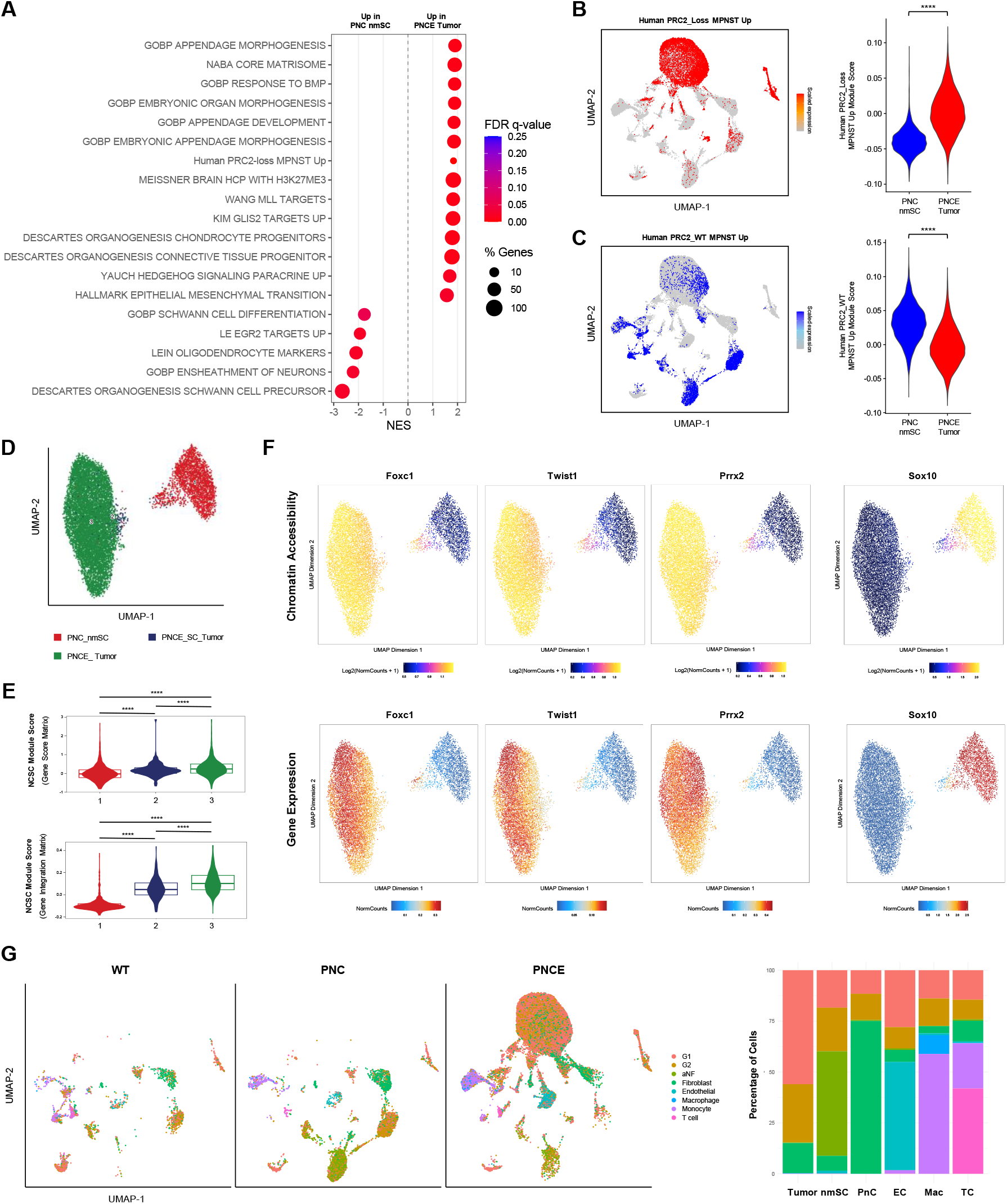
PNCE tumor cluster resembles human PRC2-loss MPNST and expresses neural crest stem cell (NCSC)-associated transcription factors. **A**. GSEA comparing PNC nmSC cluster with PNCE Tumor cluster. **B**. snRNA-seq UMAP showing module scores derived from top 300 genes upregulated in human PRC2-loss MPNSTs relative to PRC2-WT MPNSTs (left) and violin plot of the same module comparing PNC nmSC and PNCE Tumor clusters (right). Two-tailed t-test (****, p < 0.0001). **C**. snRNA-seq UMAP showing module scores derived from top 300 genes upregulated in human PRC2-WT MPNSTs relative to PRC2-loss MPNSTs (left) and violin plot of the same module comparing PNC nmSC and PNCE Tumor clusters (right). Two-tailed t-test (****, p < 0.0001). **D**. snATAC-seq UMAP of PNC nmSC cluster, PNCE SC_Tumor cluster, and PNCE Tumor cluster subset. **E**. Violin plots showing NCSC module scores from GeneScoreMatrix (chromatin accessibility) (top) and GeneIntegrationMatrix (gene expression) (bottom) across the clusters in (D). Wilcoxon rank-sum test (**** p < 0.0001). **F**. snATAC-seq UMAPs of chromatin accessibility (top) and gene expression (bottom) of representative NCSC marker genes (*Foxc1, Twist1, Prrx2*), and Schwann marker gene (*Sox10*). **G**. Projection of human MPNST single-cell dataset onto mouse sciatic nerve snRNA-seq UMAP (left) and the proportion of cells from each mouse sciatic nerve cluster mapped to corresponding clusters in the human MPNST dataset (right).

Consistent with the known role of PRC2 in H3K27 trimethylation, PNCE Tumor cluster also exhibited significant enrichment of Meissner_HCP_with_H3K27me3 genes, indicating de-repression of canonical PRC2 targets. In addition, pathways associated with epithelial to mesenchymal transition, embryogenesis, and morphogenesis were upregulated, while Schwann cell differentiation and oligodendrocyte marker pathways were suppressed. These findings indicate that PRC2 loss drives dedifferentiation and activation of developmental gene programs in *Nf1/Cdkn2a* deficient Schwann cells.

Consistent with our findings, previous studies that compare human PRC2-loss MPNST cells to PRC2-WT MPNST cells have repeatedly associated loss of PRC2 with a gain of a more dedifferentiated phenotype, particularly resembling that of neural crest stem cells (NCSCs) (26, 27). To check whether we see a similar gain of NCSC identity in the progression from PNC nmSC to PNCE Tumor, we first subset PNC nmSC, PNCE SC_Tumor, and PNCE Tumor clusters and projected the cells on the UMAP space (**Fig. 3D**). We then used a set of transcription factors (TFs) associated with NCSC identity (*Six2, Fli1, Twist1, Prrx1, Prrx2, Foxc1, Snai1, Snai2, Sox9, Otx2, Pax3, Pax6*) to construct a NCSC module to assign module scores to each cell. We found that PNCE Tumor cluster had a significantly higher NCSC module score both in terms of chromatin accessibility (GeneIntegrationMatrix) and gene expression (GeneExpressionMatrix) compared to PNC nmSC cluster (**Fig. 3E**). When visualized on the ATAC UMAP, we saw a clear upregulation of NCSC marker TFs like *Foxc1, Twist1*, and *Prrx2* in the PNCE Tumor cluster, and a concurrent downregulation of Schwann cell marker TFs like *Sox10* (**Fig. 3F**).

Lastly, to further investigate the correlation between GEMMs and human MPNST, we used published human single-cell dataset that categorized MPNSTs based on methylome and transcriptome into MPNST-G1 (group 1) and MPNST-G2 (group 2) (26). Loss of PRC2 is exclusive to MPNST-G1 with neural crest-like characteristics, while MPNST-G2 has intact PRC2 and exhibits Schwann cell precursor-like characteristics. Projection of such human MPNST datasets, including MPNST-G1, MPNST-G2, and atypical neurofibroma (aNF), onto our GEMM snRNA-seq data revealed that the majority of PNCE Tumor cluster mapped to the MPNST-G1 (56%), whereas the majority of PNC nmSC cluster corresponded to aNF (51%) (**Fig. 3G**). This suggests that *Nf1, Cdkn2a* loss models a more benign precursor lesion, while additional *Eed* deletion reprograms Schwann-lineage cells into a more malignant, dedifferentiated PRC2-loss MPNST like state.

Together, these transcriptomic and chromatin analyses demonstrate that loss of *Eed* in *Nf1/Cdkn2a*-deficient Schwann cells induces a progressive loss of Schwann identity, activation of developmental and mesenchymal transcriptional programs, and emergence of an NCSC-like dedifferentiated state that resembles human PRC2-loss MPNST.

### PNCE sciatic nerve allografts form high-grade GEM-PNSTs

To determine whether such molecular alterations can confer malignant potential, we next tested whether PRC2-loss sciatic nerves could autonomously initiate tumor formation upon transplantation. At 15 weeks post tamoxifen administration, we collected enlarged sciatic nerves from PNCE mice and transplanted small pieces of the nerve subcutaneously into immunocompetent C57BL/6 mice. After approximately a 3-month lag phase, 8 of 10 grafts developed tumors that reached the experimental endpoint of 2000mm^3 at around 5 months post graft (**Fig. 4A**). Histopathological analysis revealed densely packed, large spindle-shaped cells arranged in interlacing fascicles with a high mitotic index, features characteristic of high-grade MPNST (**Fig. 4B**). The tumors demonstrated partial loss of H3K27me3 with occasional H3K27me3-positive nuclei, likely reflecting immune cell infiltrates or *Nf1/Cdkn2a* deficient cells due to incomplete deletion of *Eed*. The tumors showed near complete loss of the Schwann cell marker S100B and strong Ki67 positivity, confirming malignant transformation and robust proliferation.

**Figure 4.**
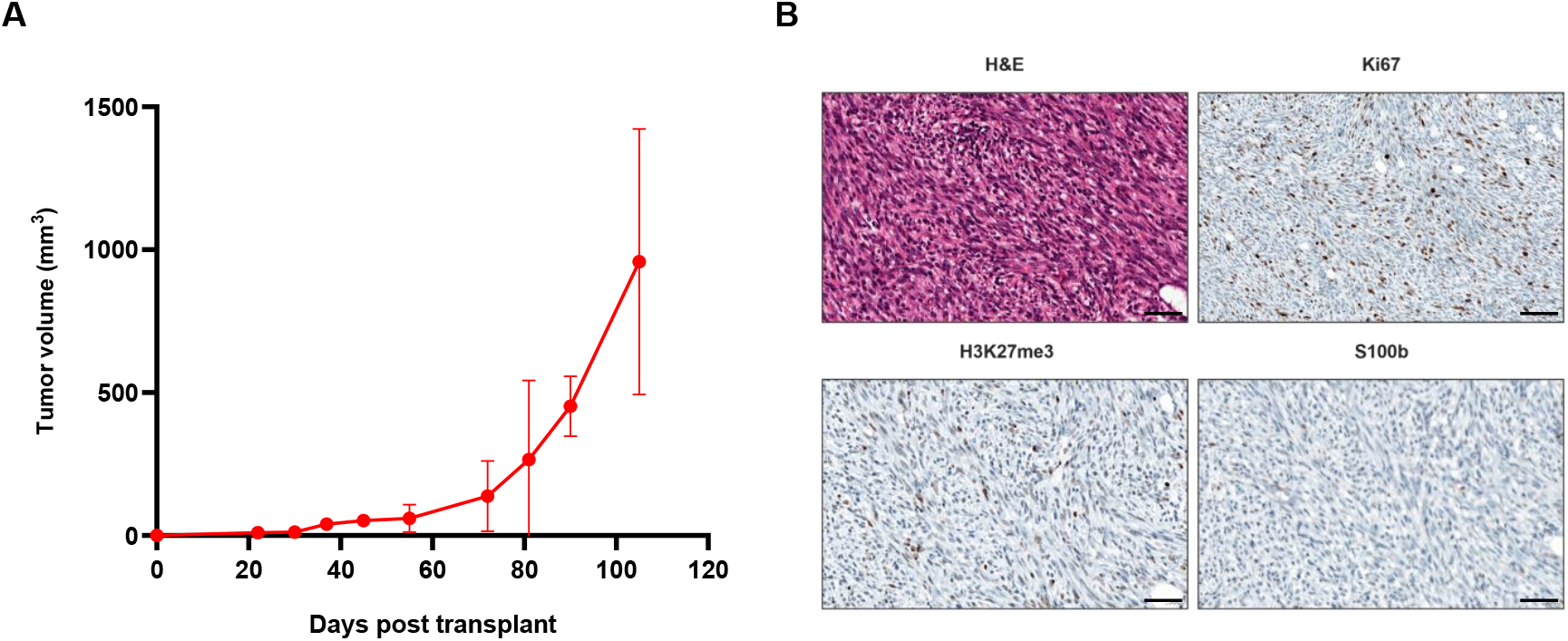
PNCE sciatic nerve allografts form high-grade GEM-PNSTs. **A**. Growth curve of PNCE sciatic nerve allografts implanted into C57BL/6J recipient mice (n = 10). Data are presented as mean ± SEM. **B**. Representative H&E and IHC staining of PNCE sciatic nerve-derived tumors. Scale bar, 50μm.

## DISCUSSION

Loss of PRC2 marks a pivotal event in the malignant transformation of MPNSTs from more benign precursors such as atypical neurofibromas. Several GEMMs have been developed that recapitulate stages of neurofibroma-to-MPNST progression. For example, deletion of *Nf1* in the Schwann cell lineage can lead to pNF development in mice (7-9), additional loss of *Cdkn2a* is associated with progression to ANF in 30-40% of mouse tumors, and ∼40% of these tumors progress to MPNST when transplanted into new mice (13, 14). Furthermore, mice that are germline heterozygous for *Nf1* and *Trp53* spontaneously develop a variety of tumors including MPNST, and further germline loss of heterozygosity reduces the time to tumor development including MPNST. In these mouse models, the MPNSTs have *Nf1*/*Trp53* and *Nf1*/*Trp53*/*Suz12* loss of heterozygosity respectively but have not been reported to develop pNF nor ANF (2, 28, 29). Despite these advances, however, a faithful model capturing the biology of PRC2 loss has been lacking.

Here, we describe a novel GEMM that recapitulates the human PRC2-loss MPNST phenotype using an inducible Plp-CreERT system to delete *Nf1, Cdkn2a*, and *Eed* in Schwann cells. This design allows temporal control of recombination and direct comparison between PRC2-intact (PNC) and PRC2-loss (PNCE) conditions. We show that the additional loss of *Eed* is sufficient to trigger rapid and diffuse tumorigenesis of the sciatic nerve, in stark contrast to the delayed, localized lesions observed in PNC mice.

Through single-nucleus RNA and ATAC Multiome analysis, we delineate how PRC2 loss reprograms Schwann lineage cells into a dedifferentiated, mesenchymal-like state. PRC2 loss disrupts Schwann cell identity through the activation of developmental and epithelial-to-mesenchymal transition (EMT)-associated programs. Importantly, we identify an intermediate population - termed the “SC_Tumor” cluster - that co-expresses Schwann and tumor markers. This hybrid state suggests an ongoing lineage transition and supports a model in which PRC2 normally constrains Schwann cells within a differentiated, lineage-committed state. Upon *Eed* deletion, this repression is lifted, enabling reactivation of early developmental transcriptional circuits. Among the transcription factors driving this reprogramming, we find neural crest stem cell (NCSC) regulators - *Foxc1, Twist1*, and *Prrx2* - showing concordant increases in both chromatin accessibility and gene expression (26, 27). This is consistent with human studies reporting that PRC2-loss MPNSTs acquire an NCSC-like, dedifferentiated phenotype within the Schwann cell lineage.

Moreover, cross-species comparisons revealed parallels between our mouse model and human MPNST. The transcriptional landscape of the PNCE tumor cluster aligned closely with human datasets representing PRC2-inactivated MPNST, whereas the PRC2-intact PNC nerves more closely resembled atypical neurofibromas. This correspondence underscores the translational relevance of our model, suggesting that PNC and PNCE mice capture distinct and sequential stages of MPNST progression.

Functionally, the PNCE sciatic nerve represents an early, lower-grade MPNST state that is captured at an earlier time point because the PNCE mice had to be euthanized early due to paralysis. The allograft setting effectively extends the observation window, demonstrating that, over time, these lesions can progress to fully transformed, high-grade MPNSTs.

In addition, the ability to study both pre-malignant (PNC) and fully transformed (PNCE) disease states in a controlled genetic background provides a powerful platform to dissect key questions in MPNST pathogenesis, such as how PRC2-loss-driven dedifferentiation alters microenvironmental interactions. It also enables direct testing of targeted therapeutic strategies, including combined treatment with MEK, CDK, and BET inhibitors.

In summary, our work describes how the loss of *Eed* in *Nf1*/*Cdkn2a*-deficient Schwann cells removes epigenetic constraints on developmental and mesenchymal programs, promoting lineage plasticity, dedifferentiation, and malignant transformation. This model not only clarifies the molecular pathogenesis of PRC2-loss MPNST but also provides an *in vivo* platform for mechanistic studies and preclinical testing of targeted therapies in this currently incurable disease.

## AUTHOR CONTRIBUTIONS

Conceptualization: **A. Patel, W. H. Cho, Y. Chen, P. Chi**

Investigation: **W. H. Cho, A. Patel, M. Khudoynazarova, S. Warda, J. Yan, D. Schoeps, E. Fishinevich, M. Pachai, C. Lee, Z. Jacobs, K. Kristoff, J. Sher**

Formal analysis: **W. H. Cho, A. Patel, D. Li**

Histology review: **C. Antonescu**

Supervision: **Y. Chen, P. Chi**

Writing: **W. H. Cho, A. Patel, Y. Chen, P. Chi**

## ACKNOWLEDGEMENTS

We thank Scott Armstrong for generously providing *Eed* flox mice respectively. The Integrated Genomics Operation (IGO) core facility at Memorial Sloan Kettering Cancer Center conducted library preparation and next-generation sequencing of samples. This work was supported in part by grants from the NIH/NCI (R01 CA228216 and DP2 CA174499), Department of Defense (W81XWH-15-1-0124 and W81XWH-22-1-0326), Francis Collins Scholar NTAP, and Cycle for Survival and Linn Family Discovery Fund to P. Chi; an NIH/NCI grant (P50 CA217694) to P. Chi and C.R. Antonescu; NIH/NCI grants (5R01CA208100-04, 5U54CA224079-03, 5P50CA092629-20) to Y. Chen; the Geoffrey Beene Cancer Research Fund to P. Chi; Translational Oncology Research in Oncology Training Program T32 grant (5T32CA160001-09) from the NIH/NCI to A.J. Patel; and an NIH grant (P30 CA008748) to MSKCC (Core Grant); IGO was funded by the NCI Cancer Center Support Grant (P30 CA08748), Cycle for Survival, and the Marie-Josée and Henry R. Kravis Center for Molecular Oncology.

---

**Figure S1.**
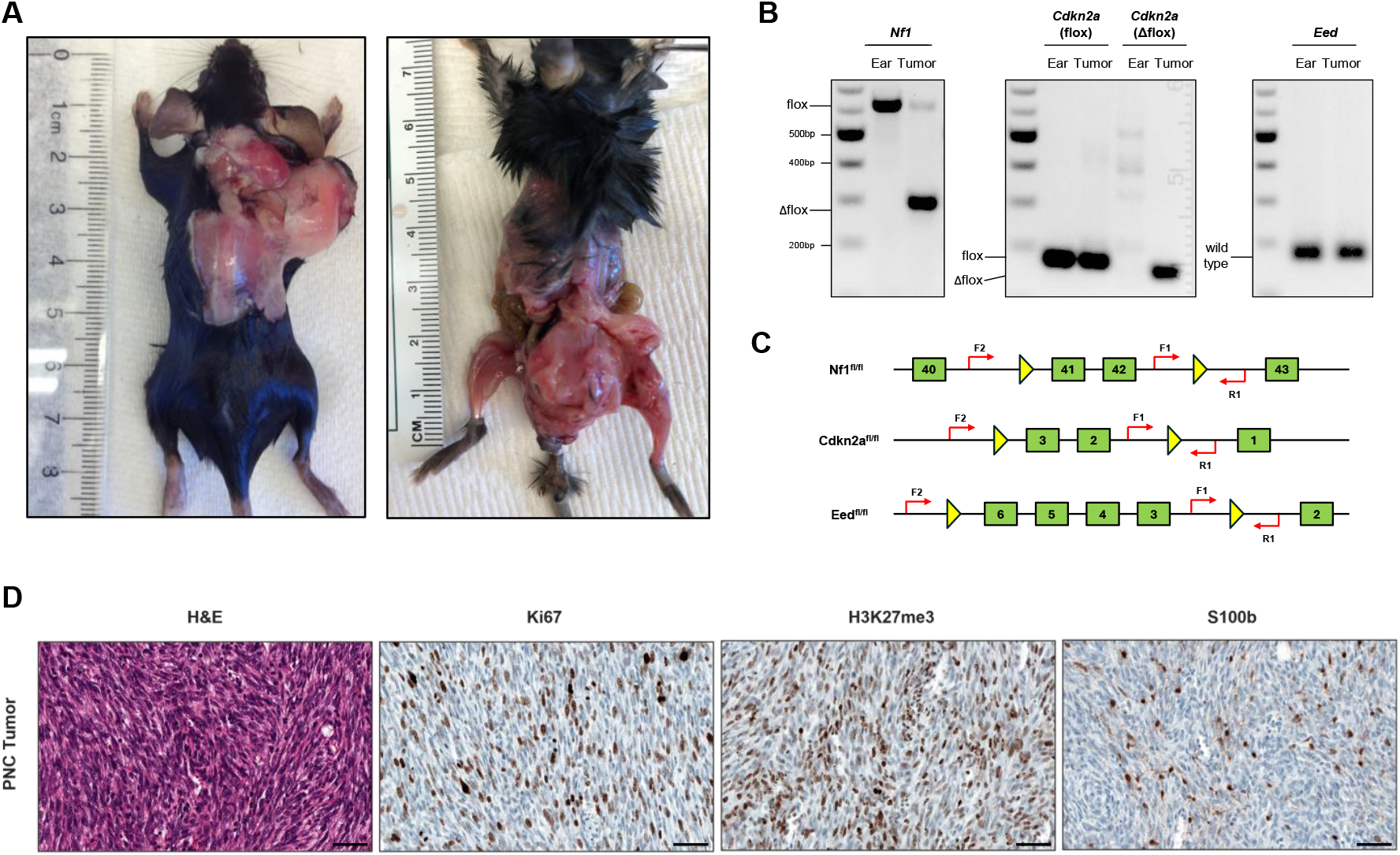
PNC mice develop tumors without sciatic nerve enlargement. **A**. Representative images of malignant tumors arising in PNC mice. **B**. PCR genotyping of PNC tumor tissue confirming Cre-mediated deletion of *Nf1* and *Cdkn2a*. **C**. Schematic of primers used for genotyping. Yellow triangle indicates loxP sites, and green boxes indicate exon positions. **D**. Representative H&E and IHC staining of PNC tumor sections collected at 104 days after neonatal tamoxifen administration. Scale bar, 50μm.

**Figure S2.**
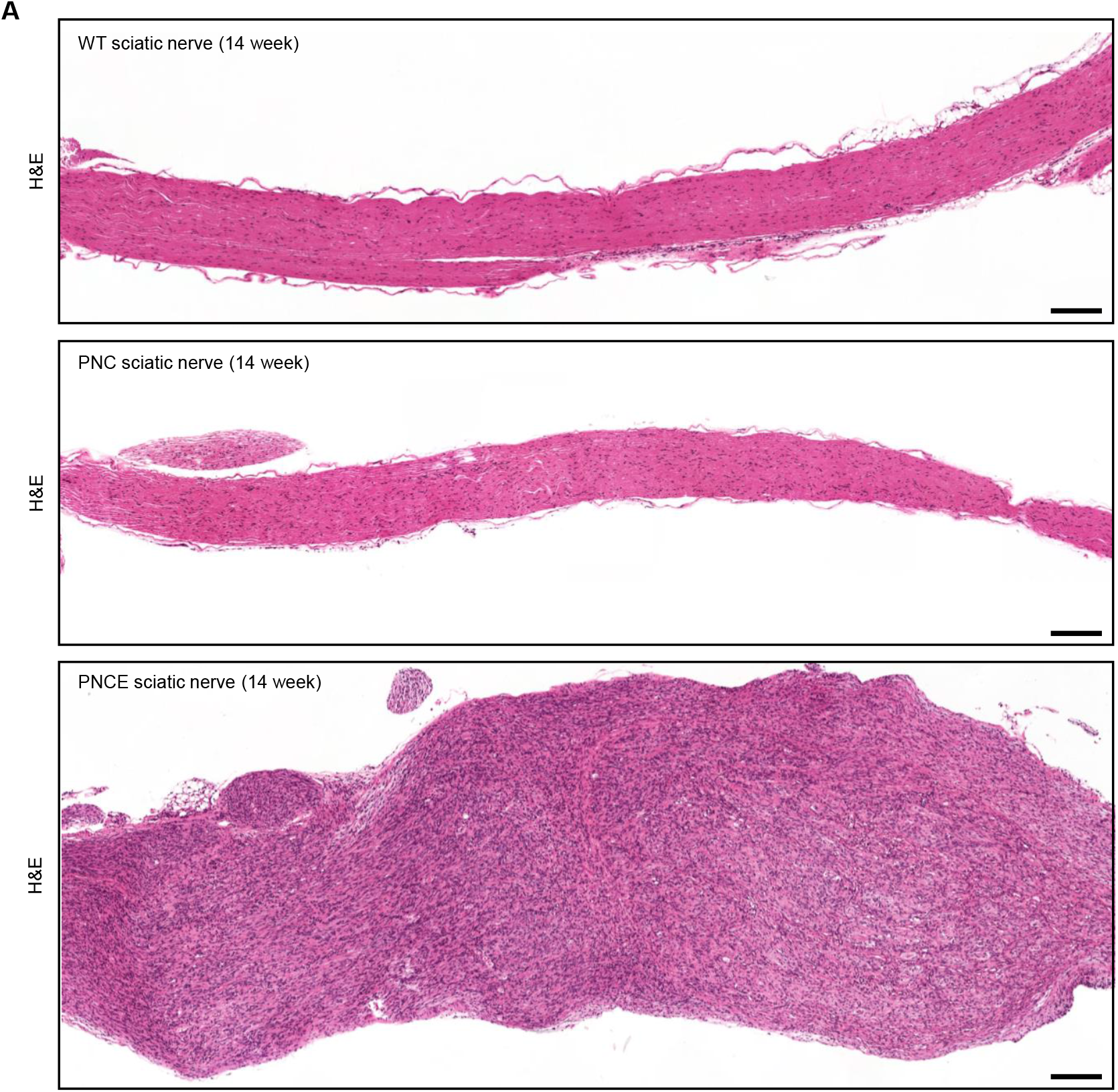
H&E staining of mouse sciatic nerve tissues. **A**. Representative H&E images of sciatic nerves from each mouse cohort at 15 weeks post tamoxifen administration. Scale bar, 200μm.

**Figure S3.**
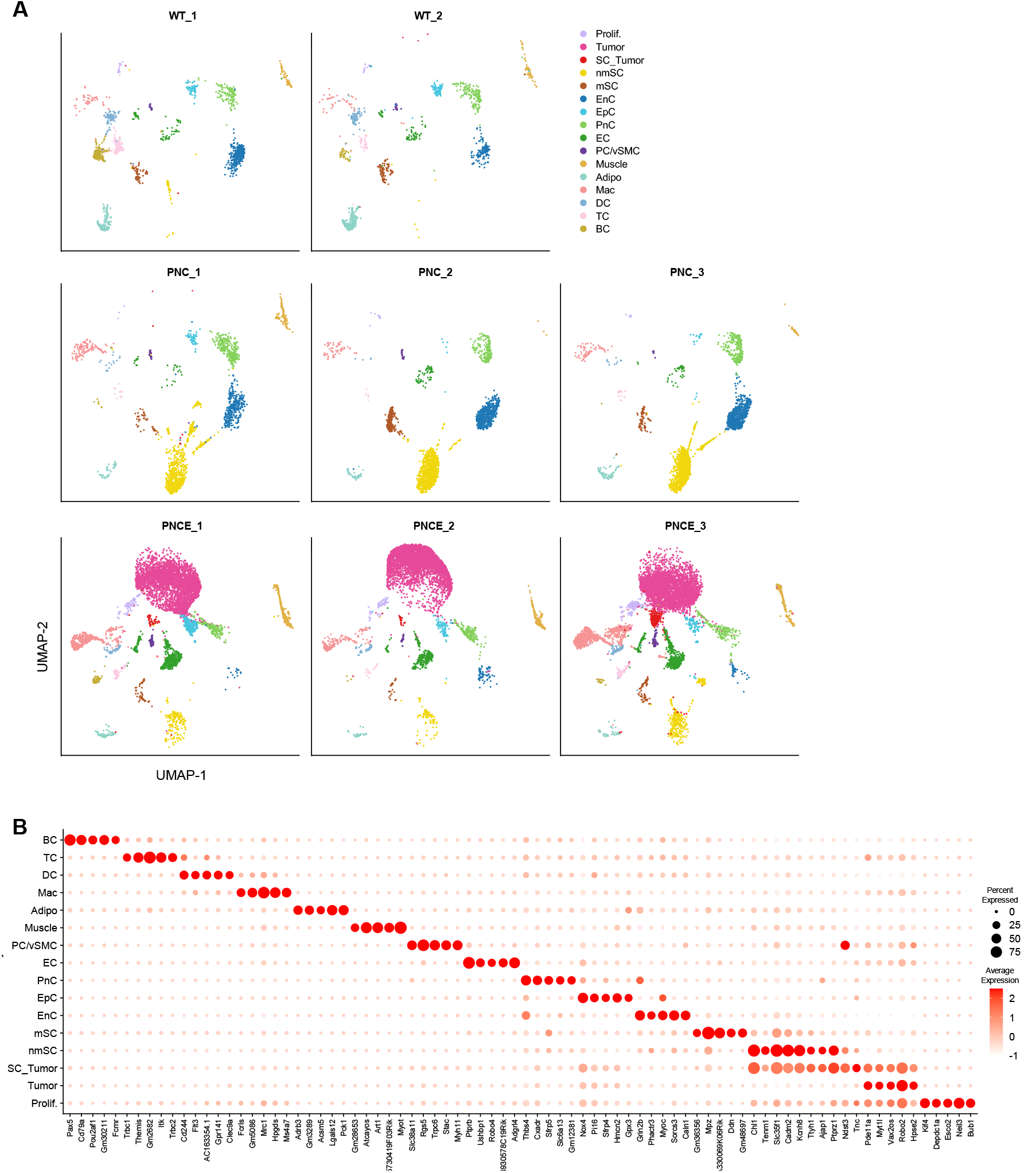
Characterization of cell clusters identified by snRNA-seq. **A**. snRNA-seq UMAPs separated by individual samples. **B**. Dot plot showing the top five marker genes defining each cluster.

